# What’s in my pot? Real-time species identification on the MinION™

**DOI:** 10.1101/030742

**Authors:** Sissel Juul, Fernando Izquierdo, Adam Hurst, Xiaoguang Dai, Amber Wright, Eugene Kulesha, Roger Pettett, Daniel J. Turner

## Abstract

Whole genome sequencing on next-generation instruments provides an unbiased way to identify the organisms present in complex metagenomic samples. However, the time-to-result can be protracted because of fixed-time sequencing runs and cumbersome bioinformatics workflows. This limits the utility of the approach in settings where rapid species identification is crucial, such as in the quality control of food-chain components, or in during an outbreak of an infectious disease. Here we present What’s in my Pot? (WIMP), a laboratory and analysis workflow in which, starting with an unprocessed sample, sequence data is generated and bacteria, viruses and fungi present in the sample are classified to subspecies and strain level in a quantitative manner, without prior knowledge of the sample composition, in approximately 3.5 hours. This workflow relies on the combination of Oxford Nanopore Technologies’ MinION™ sensing device with a real-time species identification bioinformatics application.

## Introduction

Next Generation Sequencing generates large amounts of genomic information non-specifically from low quantities of input DNA, and hence provides an unbiased way to survey the content of complex metagenomic samples with no prior knowledge ^1,2^ and without the requirement for culturing. The approach therefore has potential utility in a wide variety of contexts, such as the clinical diagnosis of infectious diseases^3^, in tracking the source of foodborne illness ^4^, monitoring crop pathogens ^5^, and environmental metagenomic studies ^6,7^. In certain contexts, such as in the diagnosis of infectious disease, the speed with which pathogenic species can be identified is of critical importance, because rapid identification enables earlier treatment^8^, and can permit the use of narrow-spectrum antibiotics which reduces the risk of bacteria developing antibiotic resistance. In an ideal world, species would be identified no more than a few hours after sample collection, which would require rapid extraction of nucleic acids, and library preparation, from the sample of interest, and minimal time spent generating and interpreting data.

Oxford Nanopore’s MinION™ is a portable nucleic acid sequencing device, which plugs directly into the USB drive of a standard laptop and requires no additional infrastructure, and is capable of generating very long reads in real time ^9^. The device contains an array of protein nanopores, in a high-salt buffer. During the sequencing reaction, an electrical potential is applied, which causes ions - and hence a current – to flow through the pores. As individual strands of DNA pass through the nanopores, the current changes in a sequence-specific manner^10^. One consequence of this is that basecalled data is generated in almost real-time: as soon as a strand passes through one of the nanopores in the array, the electrical current measurements are basecalled, and the resulting sequences are then available for downstream analysis. The time required for a strand to pass through a pore is proportional to its length at a constant translocation speed. To control the speed of strand translocation, and to ratchet the DNA strand through the pore one base at a time, the system uses an accessory protein, termed a motor. The speed of translocation can be adjusted by increasing or decreasing the concentration of ATP in the sequencing buffer. Currently, we control the speed of strand translocation to approximately ~70 bases / second as standard, meaning that a 500 base strand takes just over 7 seconds to translocate, and a 30 kb strand takes just over 7 minutes.

As each strand translocates, current measurements are written to a separate file on the laptop to which the MinION is connected. The Metrichor™ agent monitors the contents of this folder, and when a strand has finished translocating, uploads the file to the Amazon WebServer where basecalling and taxonomic classification are performed. The time taken for basecalling is proportional to the length of the strand: approximately 10 seconds for a 500 bp fragment and 10 minutes for a 30kb fragment. Classification typically takes 1-2 seconds per strand.

To complement the real-time nature of data generation, we have developed the What’s in my Pot? (WIMP) analysis pipeline on the Metrichor platform. The WIMP application classifies and identifies microbial species in real time, using a data structure that is pre-built and is shared by all application runs and which is based on taxonomy and a reference database. This data structure maps all kmers of length 24 present in the chosen database to nodes in the NCBI taxonomy tree. Because of this pre-processing, new reads can be quickly classified by looking up kmers, rather than aligning them against the original reference. During the sequencing run, the Metrichor agent uploads reads and a WIMP report is created, the read is basecalled and is classified against the prebuilt data structure. WIMP uses kraken ^11^ to map all kmers, to calculate the least common ancestor (LCA), to determine the most likely placement in the taxonomy tree, and to give each placement a classification score. This score is the fraction of all kmers found in the sequence that have been mapped to the LCA in the clade that contains the taxonomy ID. The higher this classification score, the more confident we can be that the classification was correct.

The WIMP bacterial, viral and fungal identification application works with a reference database which covers all the bacterial, viral and fungal genomes available in RefSeq ^12^. For an organism to be identified to subspecies or strain level in the WIMP report it must be present at this taxonomic level in the database. Organisms not included in the database are reported at a taxonomic group higher than strain / subspecies, such as species or genus.

To allow straightforward interpretation of WIMP results we have created an interactive report (Fig. 1), which allows the user to change several display features:

**Figure 1.**
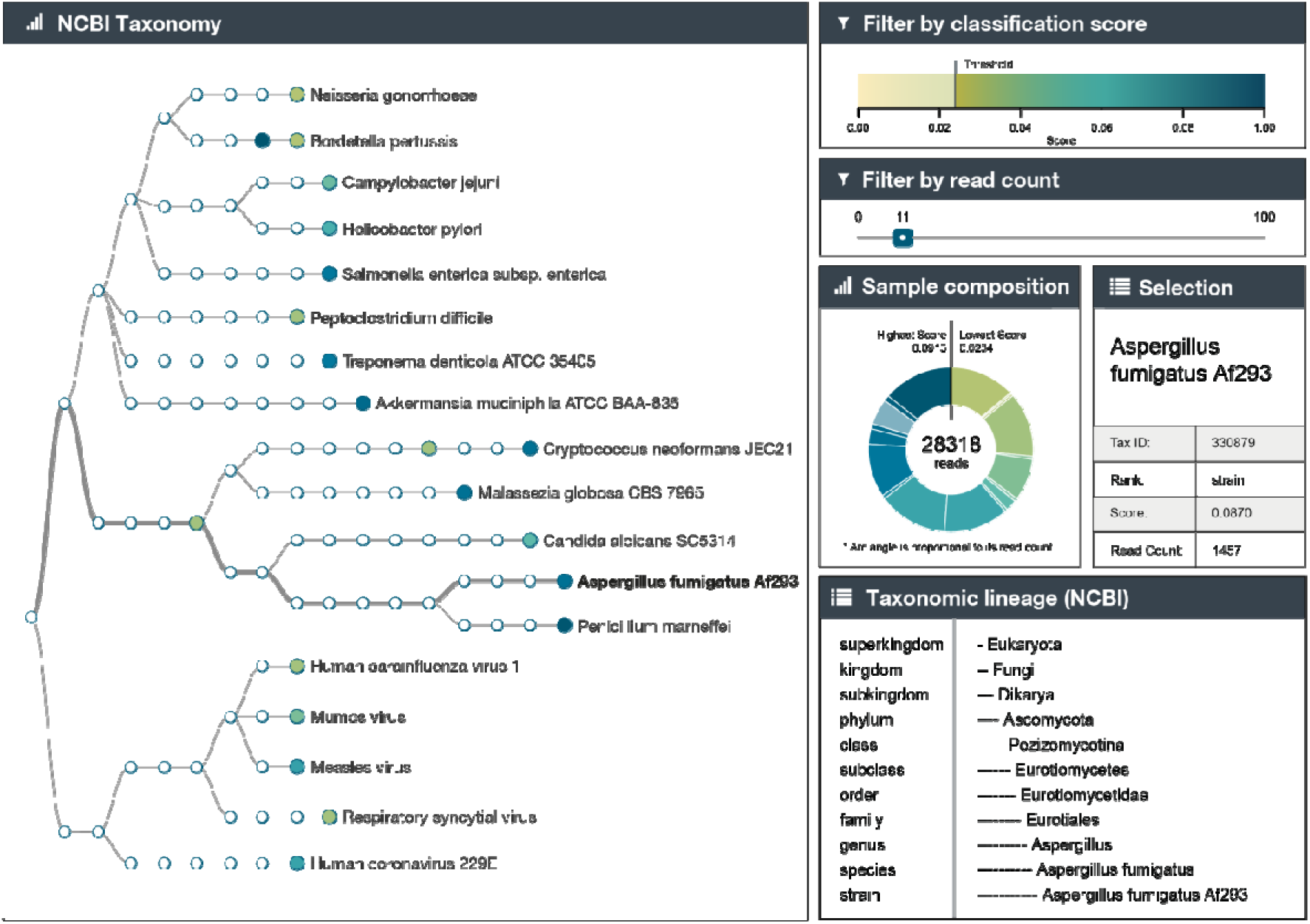
Metrichor WIMP application report, shown for bacteria, virus, and fungi.

**Sample composition:** is a donut chart showing the relative proportions of reads from the species found in the sample, and the calculated confidence level. The area of the donut segment for each species is proportional to the read count. The colour of the arc corresponds to the confidence in the **Classification score** section. Clicking on an individual segment updates the **Selection** table with the sequencing run details for that organism, shown here for Aspergillus fumigatus Af293. The selected organism also becomes highlighted in the **NCBI Taxonomy** panel (bolded text and wider lines) and is shown in a lighter colour in the donut chart.

**Selection:** gives more detail on an individual segment selected in the **Sample composition** section.

**Taxonomic lineage:** details the taxonomic classification for the individual organism selected in the **Sample composition** section.

**Classification score:** this shows the current classification threshold for displaying placements in the **NCBI Taxonomy** and **Sample composition** sections. The higher the score, the greater the confidence in the placement. The threshold can be adjusted by clicking on the coloured bar, and the charts will update automatically.

**NCBI Taxonomy:** nodes can be expanded or collapsed. Node colours correspond to those used to indicate confidence in the **Filter by classification score** and **Sample composition** sections, allowing easy interpretation of the confidence with which taxonomic calls are made.

The WIMP report is automatically updated with classified reads at regular intervals or by refreshing the browser throughout the run. An interactive version of this report is available at http://metrichor.com/workflow_instance/89822?token=U2FsdGVkXl_cDhPEc6-AkWb69BdZblg.

## Results

### What’s in my Pot?

We took several commercially-available bacterial, viral and fungal genomes, which were represented in the RefSeq database, and prepared either PCR-free genomic DNA or cDNA libraries, as appropriate. We pooled the resulting libraries together in arbitrary ratios, before sequencing, and analysis with the Metrichor application WIMP Bacteria Virus Fungi k24 for SQK-MAP006 version 1.48. All organisms were identified, down to strain or subspecies level in many cases (Fig. 1).

### Comparison of WIMP counts to concentration measured by qPCR

Next, we took six bacterial genomes and prepared low-input libraries for each, before pooling them in arbitrary ratios, and sequencing. We chose species for which published qPCR primer sequences were available, and for which the primers were robust and specific in our tests (data not shown). The sequence data was analysed in real time using the WIMP workflow. All species were successfully identified. We then compared the WIMP counts for six of these species to the relative abundance of each species in the pool as measured by qPCR. The qPCR counts agree well with the WIMP data, indicating that WIMP (including the sequencing and library preparation) is quantitative (Fig. 2). As a consequence, the limit of sensitivity of the workflow is essentially a question of how many reads are obtained from the sequencing run, which is governed by how long the device is left to generate data.

**Figure 2.**
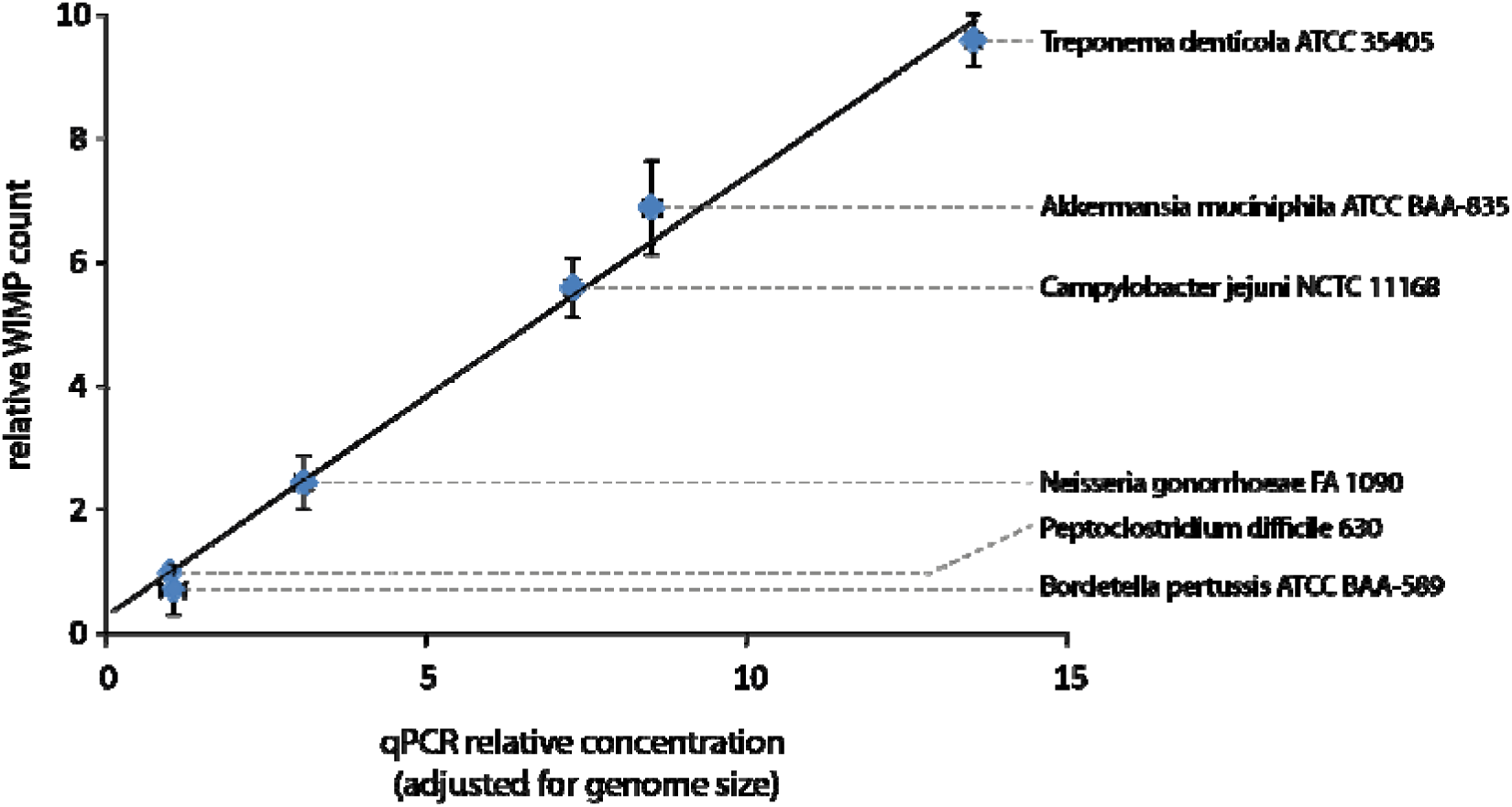
Comparison of WIMP read counts with quantitative PCR measurements

### Unpasteurised cow’s milk

We applied WIMP to a sample where we reasoned that many of the microbial species would be present in the RefSeq database: unpasteurised cow’s milk. To allow further microbial growth we left the milk samples at room temperature for two weeks before extracting the DNA, and we spiked in genomic DNA from *Listeria monocytogenes* at the earliest opportunity in the DNA extraction. The WIMP analysis reveals the presence of several bacterial genera such as *Lactococcus* and *Pseudomonas,* as well as bacteriophages, with varying degrees of confidence at different taxonomic levels (Fig. 3).

**Figure 3.**
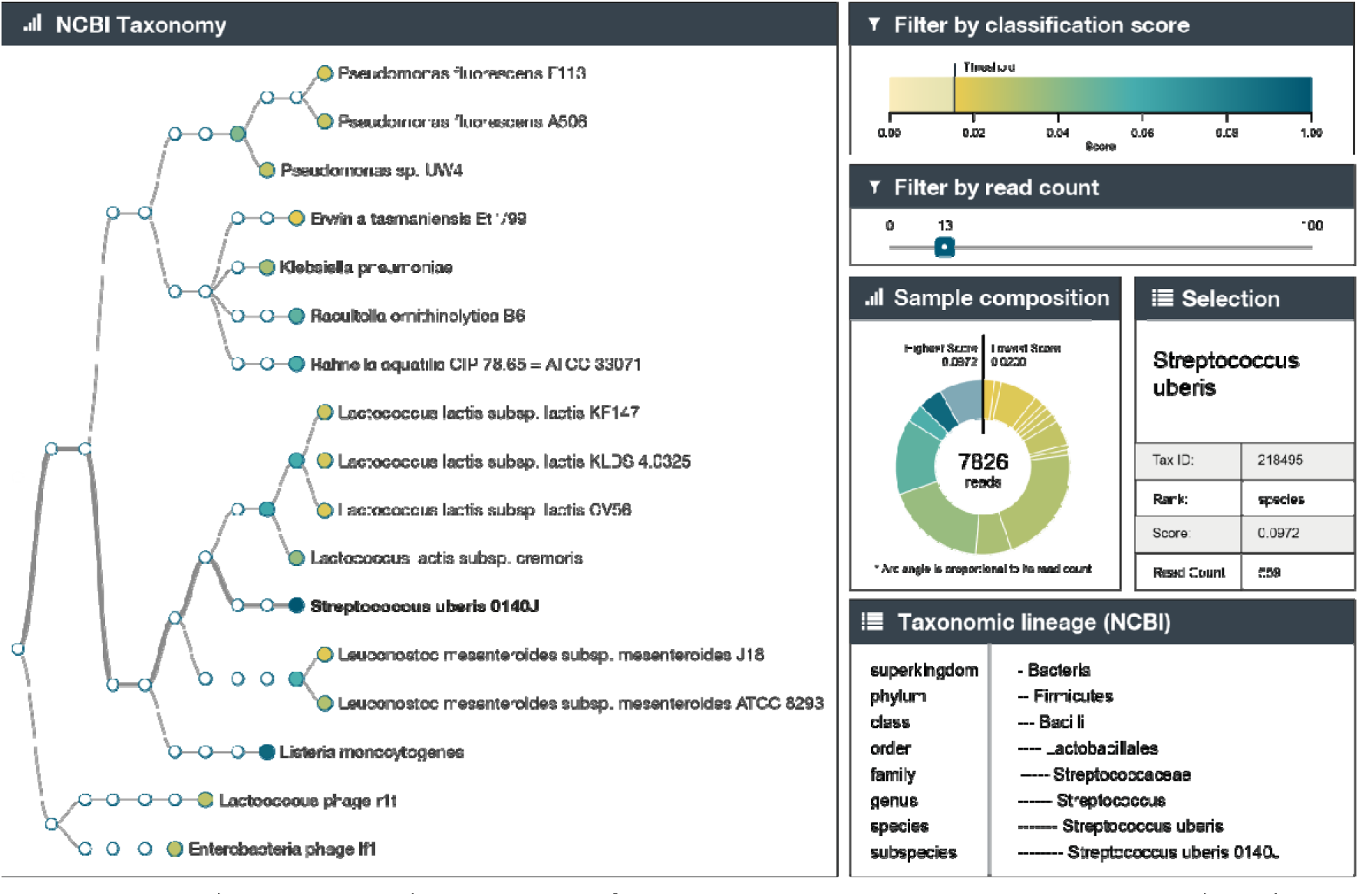
Metrichor WIMP application report for microorganisms present in unpasteurised cow’s milk (Listeria used as a spike-in).

These organisms might reasonably be expected to occur in a dairy farm environment ^13,14^. The relatively low confidence of the three strains of *Lactococcus lactis subsp lactis* indicates that none of these three strains is in the database, whereas the high confidence at the higher subspecies node indicates the presence of *Lactococcus lactis subsp lactis* albeit a different strain from those in the database. We were able to detect the spiked-in *Listeria* DNA with very high confidence, and the number of counts of listeria reads (253) matched closely the added amount, by mass (3.5% of total DNA).

We also detected bacterial species known to cause bovine mastitis: *Klebsiella pneumoniae* ^15^, and *Streptococcus uberis* ^16^ at very high confidence. Mastitis in dairy cattle is a potentially fatal disease, causing necrosis of udder tissue, and affects large proportions of cows ^17,18^. In addition to the obvious harmful consequences of udder infections in dairy cattle, these organisms can cause urinary tract infections and pneumonia in humans.

## Discussion

Using the laboratory procedure described in this manuscript, we were able to begin generation and classification of sequence data within approximately 3.5 hours starting from unpasteurised milk samples. There are several steps in this procedure that may possibly be shortened. For instance, several alternative methods are available for microbial DNA extraction from milk ^19^, some of which reportedly take less time to process samples. Additionally, to prepare double stranded DNA for sequencing on the MinION, several manipulations are currently necessary: fragment ends are blunted and phosphorylated, and a single deoxyadenosine is added to both 3’ ends. Adapters are then attached by ligation. One adapter is a hairpin, which allows both strands of the duplex to be sequenced in one read, which improves the accuracy of basecalling. The other adapter has several features which enhance the efficiency of sequencing, including a low secondary structure 5’ end for threading into the pore, and a pre-bound motor enzyme. Because there are no obligatory PCR or size-selection steps, when starting with extracted genomic DNA the entire library preparation can be performed in 90 minutes. More rapid library preparation may be performed with the use of transposases ^20^, which simultaneously shear and attach adapters to DNA fragments. In addition, it may be possible to perform the classification analysis on pre-basecalled electrical current data, by aligning this data to reference genomes which have been converted to current space using our basecalling model. In theory, such ‘squiggle-space’ analyses have the benefit of being faster to perform than base-space analyses simply because no time is spent basecalling. We intend to explore these options for possible inclusion in future versions of the WIMP application. Finally, although as of November 2015 our standard strand translocation speed is approximately 70 bases per second, we intend to release a ‘fast mode’ shortly in which strands translocate at 500 bases per second. This will allow the acquisition of data >7 times more quickly, meaning that high confidence strain identification will be possible after a shorter period of sequencing time than at present. At this translocation rate, the data presented in Fig. 1 could be obtained in under an hour.

The WIMP Metrichor application, in combination with the portable MinION sequencer is well-suited to rapidly monitoring the health and well-being of dairy animals and in performing quality control of dairy produce. With further simplification of the DNA extraction and library preparation procedures, such analyses will become possible in a non-laboratory setting, which raises the possibility of using the device in the future for applications such as the real-time monitoring of infections and their susceptibility to antibiotics.

For information on how to access the MinION and WIMP application, please visit www.nanoporetech.com

## Materials and Methods

### 1. What’s in my Pot?

Genomic DNA or RNA from *Treponema denticola* (ATCC^®^ 35405D-5™), *Akkermansia muciniphila* (ATCC^®^ BAA-835D-5™), *Campylobacter jejuni* subsp. *jejuni* (ATCC^®^ 700819D-5™), *Neisseria gonorrhoeae* (ATCC^®^ 700825D-5™), *Clostridium difficile* (ATCC^®^ BAA-1382D-5™), *Bordetella pertussis* (ATCC^®^ BAA-589D-5™), *Salmonella enterica subsp. enterica* serovar Typhimurium (ATCC^®^ 700720D-5™), *Helicobacter pylori* (ATCC^®^ 43504D-5™), Human respiratory syncytial virus (ATCC^®^ VR-955D™), Human coronavirus 229E (ATCC^®^ VR-740D™), Measles virus, strain Edmonston (ATCC^®^ VR-24D™), Mumps virus, strain Enders (ATCC^®^ VR-106D™), Human parainfluenza virus 1 (ATCC^®^ VR-94D™), *Aspergillus fumigatus* Fresenius (Af293, ATCC^®^ MYA-4609D-2™), *Candida albicans* (strain SC5314, ATCC^®^ MYA-2876D-5™), *Cryptococcus neoformans* (JEC21, ATCC^®^ MYA-565D-5™), *Malassezia globosa* (CBS 7966, ATCC^®^ MYA-4612D-5™), and *Penicillium marneffei* (QM 7333, ATCC^®^ 18224D-2™) were all purchased from ATCC.

The bacterial and fungal genomic DNAs were sheared to ~10kb in a g-TUBE (Covaris) according to the manufacturer’s protocol.

The viral genomic RNAs were reverse-transcribed and PCR-amplified using SuperScript^®^ III One-Step RT-PCR System (Thermo Fisher) and the appropriate primers (Table 1):

**Table 1.**
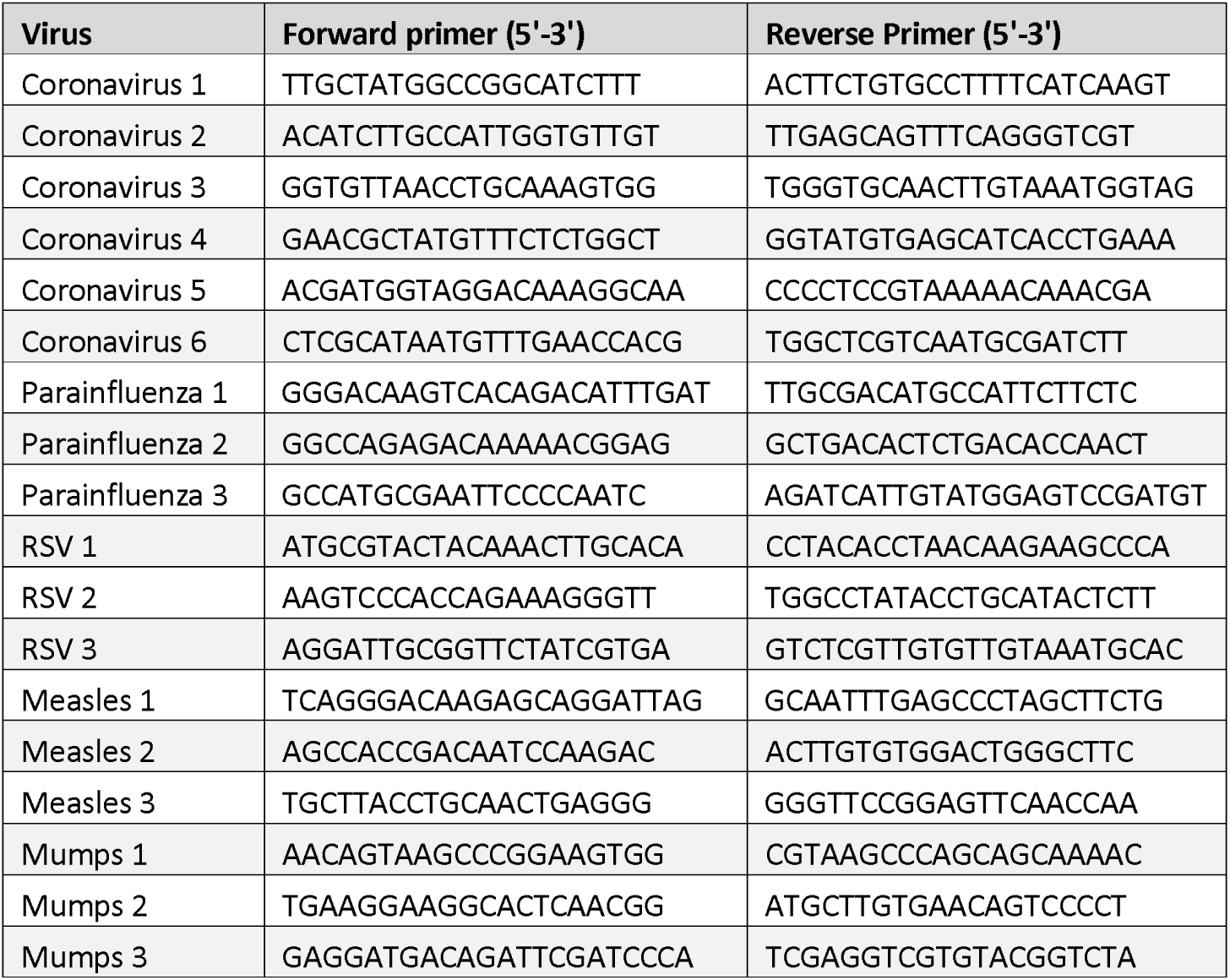
Primers used in viral reverse transcription

Bacterial and fungal genomic DNAs and viral cDNA amplicons were end-repaired and dA-tailed using NEBNext^®^ Ultra™ II End Repair/dA-Tailing Module (New England Biolabs). We prepared sequencing libraries using Oxford Nanopore Technologies’ version 6 library preparation kit, following Oxford Nanopore’s protocols, and sequenced for 6 hours following Oxford Nanopore’s standard running protocols. The samples were analysed with Metrichor application WIMP Bacteria Virus Fungi k24 for SQK-MAP006 version 1.48 as described in the text.

### 2. Comparison of WIMP counts to concentration measured by qPCR

Genomic DNA from each of *Treponema denticola, Akkermansia muciniphila, Campylobacter jejuni* subsp. *jejuni, Neisseria, Clostridium difficile,* and *Bordetella pertussis* were sheared and end-prepared as described above, and Oxford Nanopore’s low-input adapters were ligated on. The libraries were PCR-amplified according to Oxford Nanopore’s low-input PCR protocol. The PCR product was purified using 0.5x volume of Agencourt AMPure XP beads (Beckman Coulter) and the eluted DNA was used as template for qPCR and for preparing sequencing libraries as described above. The samples were analysed with Metrichor application WIMP Bacteria k24 for SQK-MAP006 version 1.34.

qPCR standard curves were made in triplicate for all bacteria individually using the primers listed in Table 2:

**Table 2.**
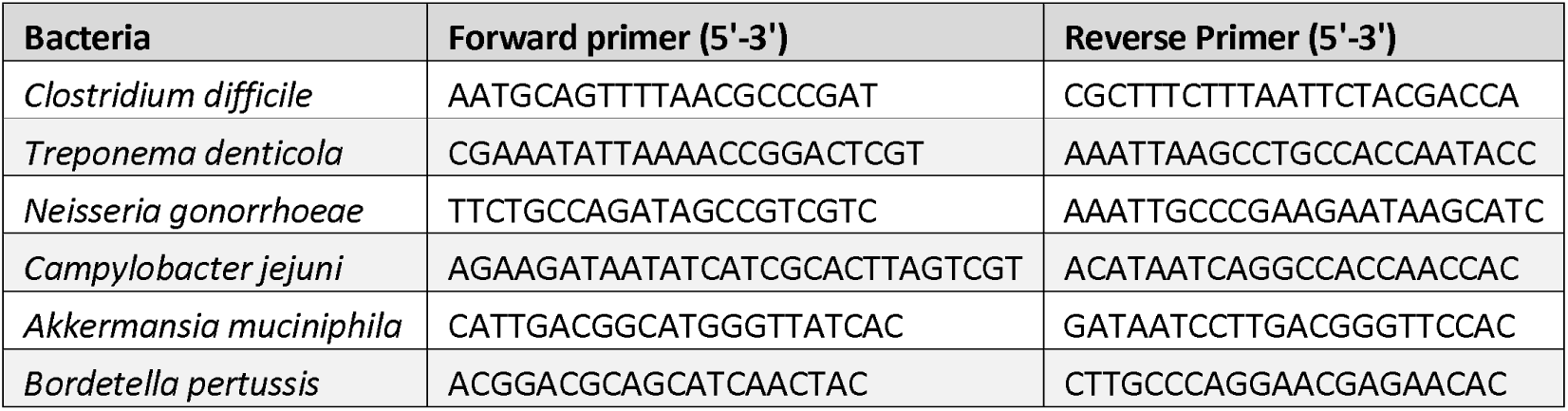
qPCR primers for bacteria

10 ng of pooled bacterial DNA was used in each of three replicates for each primer pair to determine the concentration of each bacterial species in the template mix. When a standard curve is used, qPCR measures the number of copies of a bacterial genome, whereas at a constant fragment size, WIMP counts are proportional to the genome size, the qPCR results were adjusted for genome size before being normalised to *Clostridium difficile,* which had the fewest WIMP reads.

### 3. Unpasteurised milk

Raw cow’s milk was purchased from Udder Milk Creamery Co-Op (http://www.uddermilk.com/) and immediately aliquoted into sterilized plastic tubes or glass flasks. Aliquots were left at room temperature for two weeks to allow microbial growth. We extracted genomic DNA from 4 mL of raw milk using the Milk Bacterial DNA Isolation Kit (Norgen Biotek Corp). We followed the manufacturer’s protocol with the addition of an initial centrifugation of the milk at 1,000 rpm for 2 minutes to pellet the host cow cells. The supernatant from this centrifugation included the bacterial cells and was transferred to a new tube. The bacterial cells were pelleted by centrifuging at 14,000 rpm for 3 min. 25 ng corresponding to roughly 3.5% of the total DNA, *Listeria monocytogenes* (Strain EGDe ATCC^®^ BAA-679D-5™) was added to the sample before binding of DNA to the column.

We prepared PCR-free sequencing libraries using Oxford Nanopore Technologies’ version 6 library preparation kits, following Oxford Nanopore’s protocols, and sequenced the samples with and without spiked-in *Listeria* on different flow cells, for 3 hours each, following Oxford Nanopore’s standard running protocols. The samples were analysed with Metrichor application WIMP Bacteria Virus Fungi k24 for SQK-MAP006 version 1.48.

